# Pyruvate kinase controls signal strength in the insulin secretory pathway

**DOI:** 10.1101/2020.01.15.907790

**Authors:** Sophie L. Lewandowski, Rebecca L. Cardone, Hannah R. Foster, Thuong Ho, Evgeniy Potapenko, Chetan Poudel, Halena R. VanDeusen, Tiago C. Alves, Xiaojian Zhao, Megan E. Capozzi, Ishrat Jahan, Craig S. Nunemaker, Jonathan E. Campbell, Craig J. Thomas, Richard G. Kibbey, Matthew J. Merrins

## Abstract

Pancreatic β-cells couple nutrient metabolism with appropriate insulin secretion. Here, we show that pyruvate kinase (PK), which converts ADP and phosphoenolpyruvate (PEP) into ATP and pyruvate, underlies β-cell sensing of both glycolytic and mitochondrial fuels. PK present at the plasma membrane is sufficient to close K_ATP_ channels and initiate calcium influx. Small-molecule PK activators increase β-cell oscillation frequency and potently amplify insulin secretion. By cyclically depriving mitochondria of ADP, PK restricts oxidative phosphorylation in favor of the mitochondrial PEP cycle with no net impact on glucose oxidation. Our findings support a compartmentalized model of β-cell metabolism in which PK locally generates the ATP/ADP threshold required for insulin secretion, and identify a potential therapeutic route for diabetes based on PK activation that would not be predicted by the β-cell consensus model.

**GRAPHICAL ABSTRACT:** 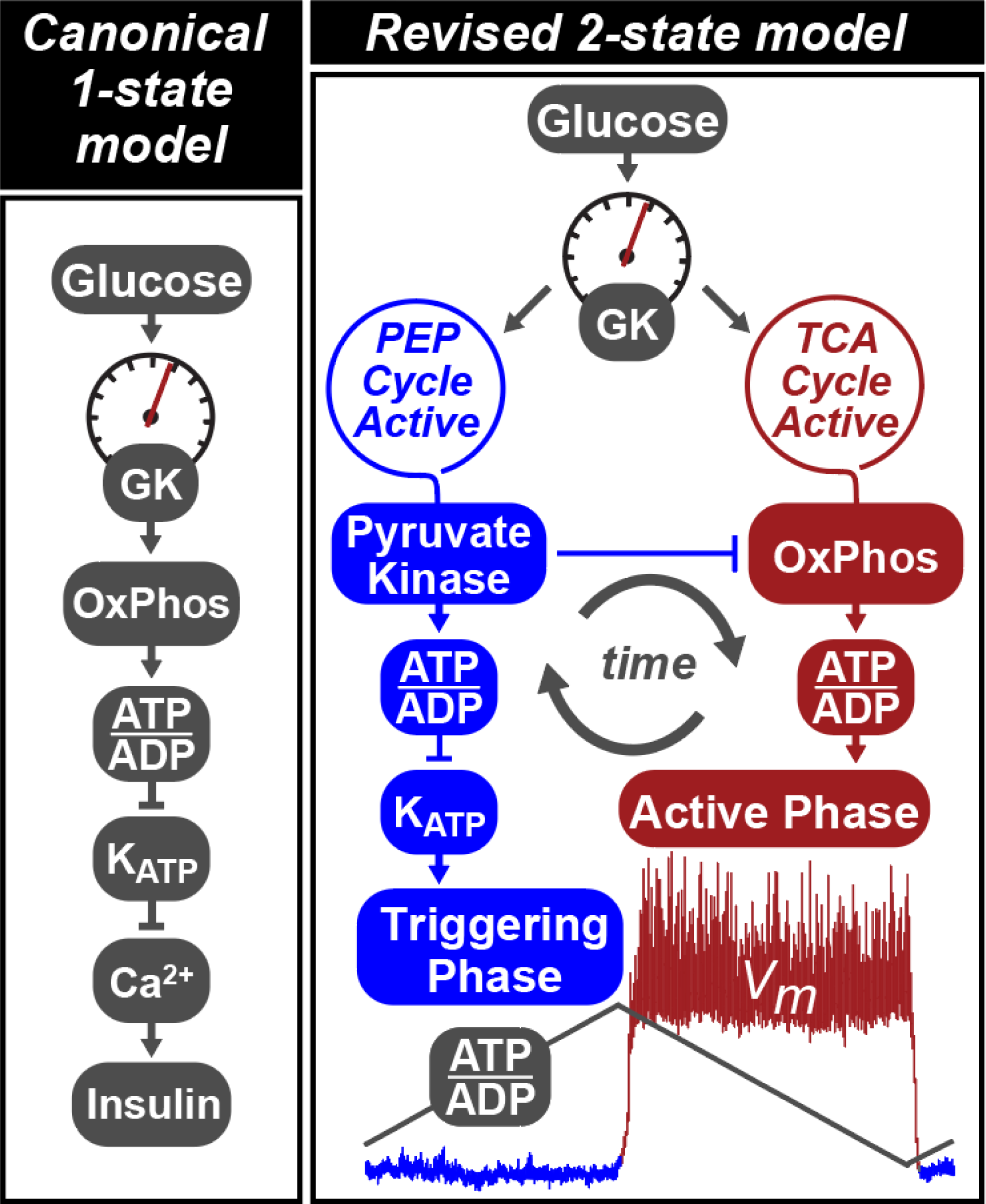

The consensus model for β-cell glucose sensing supports a dominant role for OxPhos. This model doesn’t fully explain the observed metabolic and electrophysiologic oscillations associated with glucose-stimulated insulin secretion. Lewandowski *et al*. challenge this model by mechanistically connecting the anaplerotic PEP cycle to the electrically silent triggering phase, and OxPhos to the electrically active secretory phase. Here, the allosteric recruitment of pyruvate kinase directs metabolic traffic between the two cycles and identifies potential therapeutic strategies for diabetes based on pharmacologic pyruvate kinase activation.

**HIGHLIGHTS:**

- Compartmentalized pyruvate kinase (PK) activity underlies β-cell fuel sensing
- Membrane-associated PK closes K_ATP_ channels and controls calcium influx
- By lowering ADP, PK toggles mitochondria between OxPhos and PEP biosynthesis
- Pharmacologic PK activation increases oscillatory frequency and amplifies secretion

## INTRODUCTION

A characteristic feature of pancreatic β-cells is their ability to couple metabolic glucose sensing with appropriate insulin secretion. The most widely-accepted description of the sensing mechanism involves the oxidation of glucose carbons in the mitochondria to generate a proton motive force that, through ATP synthase, sequentially raises the ATP/ADP ratio, closes K_ATP_ channels, and activates Ca^2+^ influx, which triggers insulin granule fusion with the plasma membrane (Prentki et al., 2013). Many different components of the glucose-sensing apparatus are well-characterized. For example, glucokinase (GK) and K_ATP_ channels are genetically linked to insulin secretion through both gain and loss of function mutations in humans (Nichols, 2006). However, several lines of evidence challenge one key component of the canonical mechanism – the exclusivity of coupling oxidative phosphorylation (OxPhos) to K_ATP_ channel closure.

A central tenet of mitochondrial respiratory control is that, in the presence of adequate O_2_ and substrate, mitochondrial OxPhos is dependent on ADP availability (Chance and Williams, 1955). Workload in the form of ATP hydrolysis is therefore the principal driver for OxPhos in many cells. Although the β-cell ATP/ADP ratio is unusually substrate sensitive, a work dominated drive for OxPhos poses a challenge to the canonical β-cell model, since ADP privation is the physiological driver of K_ATP_ channel closure (Koster et al., 2005; Nicholls, 2016). Consistent with ADP limitation, mitochondrial respiration is highest after membrane depolarization, rather than during the triggering phase when K_ATP_ channels close (Jung et al., 2000). Furthermore, calcium influx precedes oxygen consumption during glucose-stimulated oscillations (Kennedy et al., 2002), implying that the canonical model does not hold at steady state.

If the dependence of K_ATP_ closure on OxPhos is to be questioned, is there an alternative ATP/ADP generator that is limited by glucose and functions prior to membrane depolarization? One clue may be that anaplerotic flux through pyruvate carboxylase (PC) is more strongly correlated with insulin secretion than oxidative flux through pyruvate dehydrogenase (PDH) (Alves et al., 2015; Fransson et al., 2006; MacDonald et al., 2005; Prentki et al., 2013). Glucose carbons that transit through PC generate 40% of cytosolic PEP through the cataplerotic mitochondrial PEP carboxykinase (PEPCK-M) reaction (Stark et al., 2009). This ‘PEP cycle’ has been linked to insulin secretion (Jesinkey et al., 2019; Stark et al., 2009) and provides a mechanism distinct from OxPhos for cytosolic ATP/ADP generation via pyruvate kinase (PK), which is allosterically activated by fructose 1,6-bisphosphate (FBP) prior to membrane depolarization (Merrins et al., 2013, 2016).

Here we provide evidence for a revised model of β-cell metabolism based on the ability of PK to initiate K_ATP_ channel closure at the plasma membrane. In this model, cytosolic ADP lowering by PK-driven PEP hydrolysis deprives mitochondria of ADP, at the same time creating antiphase OxPhos oscillations. Rather than triggering depolarization, mitochondrial OxPhos provides the energy to sustain membrane depolarization and insulin secretion. Through a mechanism that would not be predicted by the canonical pathway, pharmacologic activation of PK amplifies the metabolic response without increasing glucose oxidation, providing a potential new therapeutic strategy for diabetes that does not inappropriately trigger insulin secretion at low glucose. In a companion paper in this issue (Abulizi et al., 2019), we show that the control of glucose signal strength by PK is dependent on PEPCK-M, and demonstrate the *in vivo* relevance of small molecule PK activation as a therapeutic strategy in pre-clinical models of diabetes.

## RESULTS

### Membrane-associated pyruvate kinase is sufficient to close K_ATP_ channels

The premise for local control of K_ATP_ channel closure by glycolysis is established in cardiac myocytes (Weiss and Lamp, 1987; Dhar-Chowdhury et al., 2005; Hong et al., 2011). To test this concept in mouse β-cells, K_ATP_ channels were recorded in the inside-out patch clamp configuration to investigate their relationship with membrane-associated PK (**Figure 1A**). K_ATP_ channel opening occurred spontaneously after patch excision, and the channels were reversibly inhibited by 1 mM ATP. A pretest solution containing the K_ATP_-opener ADP (0.5 mM) restored channel activity. When applied to the ADP-containing solution, PEP reduced K_ATP_ current or closed the channels completely (**Figure 1B**), indicating the presence of PK associated with the plasma membrane in mouse islets. PEP reduced the average K_ATP_ channel power (reflecting the total number of transported ions) by 80% (**Figure 1C**), which was due to a 73% decrease in the frequency of events (**Figure 1D**), as well as a 68% decrease in the fractional time in the open state (**Figure 1E**). We observed no significant effect of ADP and PEP on current amplitude (**Figure 1F**). In K_ATP_ channels from dispersed human islets, similar trends were observed on channel power, frequency, open time, and amplitude (**Figures 1G-K, S1**, and **Table S1**). These data indicate that, even in the presence of high ADP in the bath, endogenous PK is able to locally raise the ATP/ADP ratio to close K_ATP_ channels.

**Figure 1.**
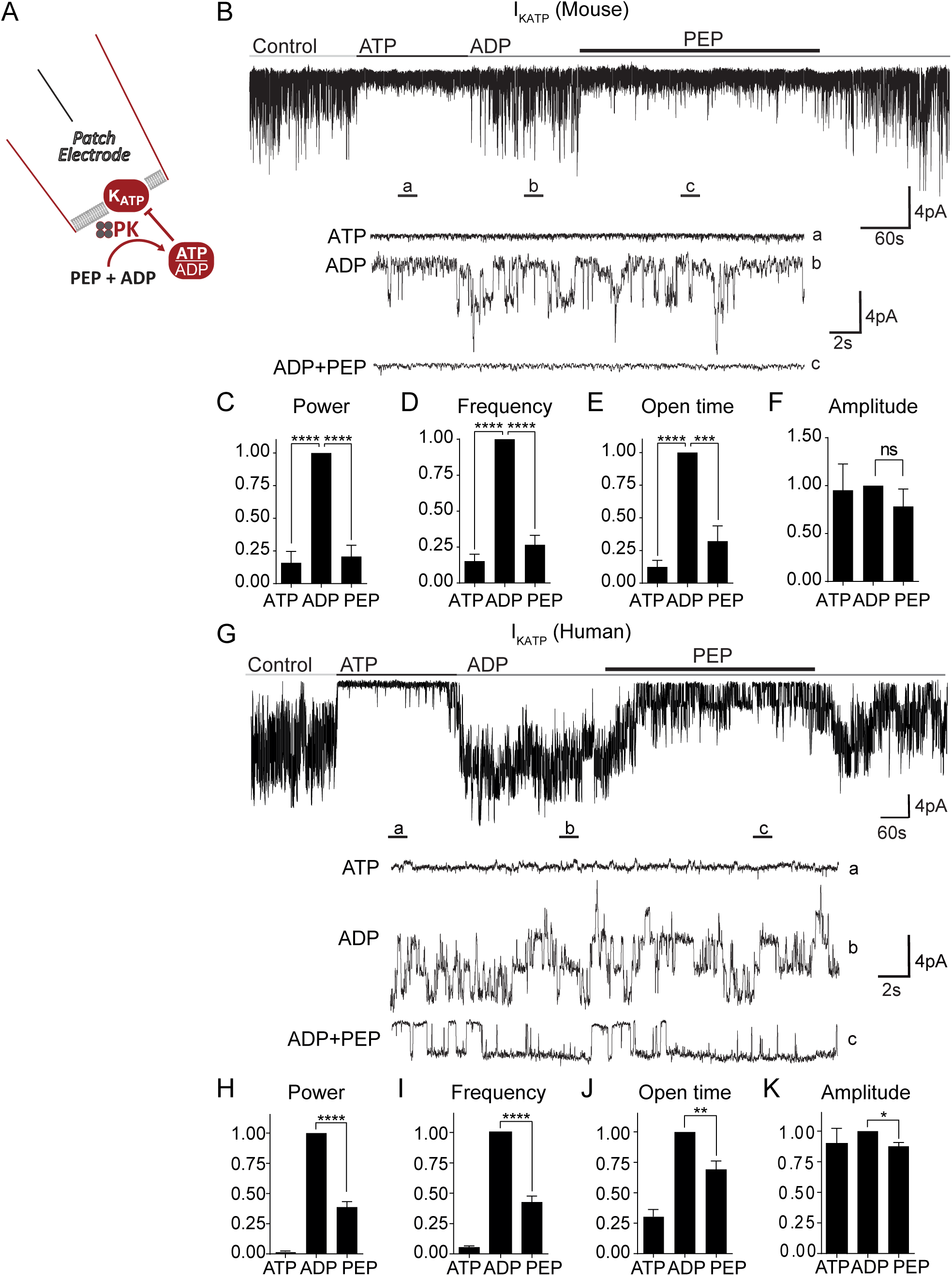
Membrane-associated PK is sufficient to close K_ATP_ channels. (A) Experimental setup for excised patch of K_ATP_ channels. (B) Applying the substrates for PK closes K_ATP_ channels in mouse islets. ATP, 1 mM ATP; ADP, 0.5 mM ADP + 0.1 mM ATP; PEP, 5 mM phosphoenolpyruvate. Holding potential, −50 mV. (C-F) Analysis of K_ATP_ channel closure in terms of power (C), frequency (D), open time (E), and amplitude (F) in mouse islets. (G) Applying the substrates for PK closes K_ATP_ channels in human islets. (H-K) Analysis of K_ATP_ channel closure in terms of power (H), frequency (I), open time (J), and amplitude (K) from 3 human islet donors. Data are shown as mean ± SEM. *p < 0.05, **p < 0.01, ***p < 0.001, ****p < 0.0001 by 1-way ANOVA. See also Figure S1 and Table S1.

### PK recruitment potentiates insulin secretion from rodent and human islets

Since GK determines the entry of carbons into glycolysis, the consensus model predicts that activating distal glycolysis should have no impact on secretion. Nevertheless, given the relationship of PK to K_ATP_ channels (**Figure 1**), we took advantage of the small-molecule PK activator (PKa) TEPP-46 to stabilize the active forms of recruitable PK (Anastasiou et al., 2012). Pancreatic β-cells express three isoforms of PK (M1, M2, and L) (DiGruccio et al., 2016; Mitok et al., 2018) (**Figure 2A**). While PKM1 is stably active, the recruitable PKM2 and PKL isoforms are allosterically activated by glycolytic FBP. PKa increased the activity of recombinant PKM2 and PKL, while having no effect on PKM1 (**Figure 2B**). Acute application of PKa (10 μM) to INS1 832/13 cells significantly increased PK activity in cell lysates (**Figure 2C**), and augmented insulin secretion (**Figure 2D**). In mouse islets, PKa increased insulin secretion 4-fold by comparison to control (**Figure 2E**), but did not significantly impact simultaneously measured glucagon secretion (**Figure 2F**). To control for off-target effects, we tested several alternative PK activators with different chemical scaffolds (Boxer et al., 2010; Jiang et al., 2010; Anastasiou et al., 2012). Six PK activators were capable of enhancing insulin secretion and one was not (**Figure S2**); although TEPP-46 (referred to as PKa in this manuscript) was not the most effective secretagogue, it is commercially available and was chosen for reproducibility.

**Figure 2.**
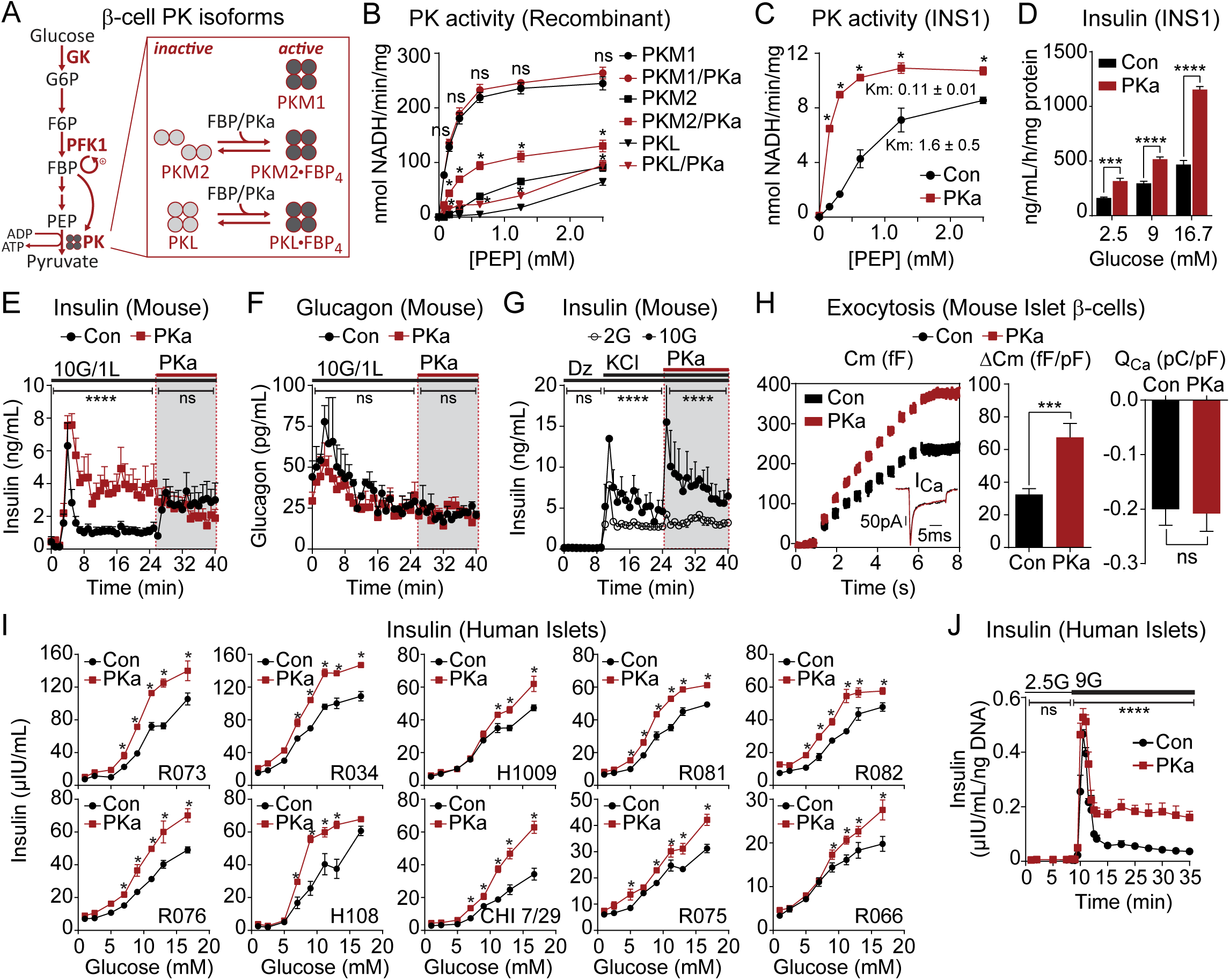
Pharmacological activation of β-cell PK enhances glucose-stimulated insulin secretion (GSIS) from rodent and human islets. (A) Schematic depicting PKM2 and PKL activation by glycolytic FBP or PKa. (B) PKa (10 µM TEPP-46) increased the activity of recombinant PKM2 and PKL with no effect on PKM1 (n = 3). (C) PKa increased PK activity in INS1 832/13 lysates (n = 3). (D) PKa increased GSIS from INS1 832/13 cells in static incubation assays (n = 6). (E-F) PKa increased GSIS from perifused mouse islets when applied with 10 mM glucose (10G) or during second phase (shaded box) (E) with no concurrent effect on glucagon secretion (F) (n = 3). (G) PKa increased KCl-stimulated insulin secretion only in the presence of high glucose (n = 3). (H) PKa increased insulin exocytosis from mouse islet β-cells when applied via patch pipette without affecting calcium current (I_Ca_) or calcium influx (Q_Ca_) (n = 20 cells per treatment). (I) PKa increased GSIS from human donor islets in static incubation assays. Data points represent the mean of 4 technical replicates for each experiment. (J) PKa increased insulin secretion from human islets (donor H108) perifused with 9 mM glucose. (9G) but not 2.5 mM glucose (2.5G). Data points represent the mean of 4 technical replicates for each experiment. Data are shown as mean ± SEM. *p < 0.05, ***p < 0.001, ****p < 0.0001 by 1-way ANOVA (B-D, I), 2-way ANOVA (E-G, J), or t-test (H). See also Figures S2-3 and Table S1.

PK activation was similarly effective at augmenting depolarization-induced exocytosis. In the control condition, the application of 30 mM KCl (in the presence of diazoxide) elicited biphasic insulin secretion that was more pronounced at elevated glucose (**Figure 2G**). The subsequent addition of PKa re-initiated biphasic release and further elevated secretion when glucose was present at 10 mM, but not at 2 mM, indicating that the enhancement of secretion by PKa is fuel dependent. When applied to islet β-cells via patch pipette, PKa increased exocytosis by ∼110% in response to 10 step depolarizations, without any change in the evoked calcium current, confirming that PK also can amplify insulin secretion independently of K_ATP_ channel closure (**Figure 2H**).

To determine if the effect of PK activation in rodent tissues is relevant to humans, we tested the insulin secretory response to glucose (between 1 and 16.7 mM) in re-aggregated islets from 10 human donors across a spectrum of age and BMI (**Figure 2I** and **Table S1**). Insulin secretion was significantly amplified by PKa at glucose concentrations >5 mM, without significant changes at subthreshold glucose concentrations or in the half-maximal effective concentration (EC50) of glucose (vehicle, 10.8 ± 2.5 mM; PKa, 9.1 ± 1.1 mM, *n* = 10, *P* > 0.05) (**Figure S3**). In a dynamic perifusion assay, PKa had no effect on basal or first-phase insulin secretion, and increased second-phase secretion approximately 3.5-fold (**Figure 2J**). As in mouse islets, these data demonstrate that the actions of PK in human islets are dependent on fuel availability.

### PK exerts GK-independent effects on the β-cell triggering and amplifying pathways

GK lacks feedback inhibition and is responsive to glucose across the physiologic range and, consequently, determines the rate of β-cell glucose metabolism (Matschinsky and Ellerman, 1968). Despite that PK is sufficient to close K_ATP_ (**Figure 1**), the lack of a threshold effect makes it unlikely that PK activation enhances insulin secretion by increasing fuel intake (**Figure 2**). To test this idea further, PK activation was compared to GK activation in human islets (**Figure 3A**). PKa primarily increased the amplitude of insulin secretion with only a small effect on the EC50, while GK activation reduced the EC50 for glucose with a lesser effect on the amplitude. When combined, the effects of PK and GK activation were additive, indicating separable mechanisms.

**Figure 3.**
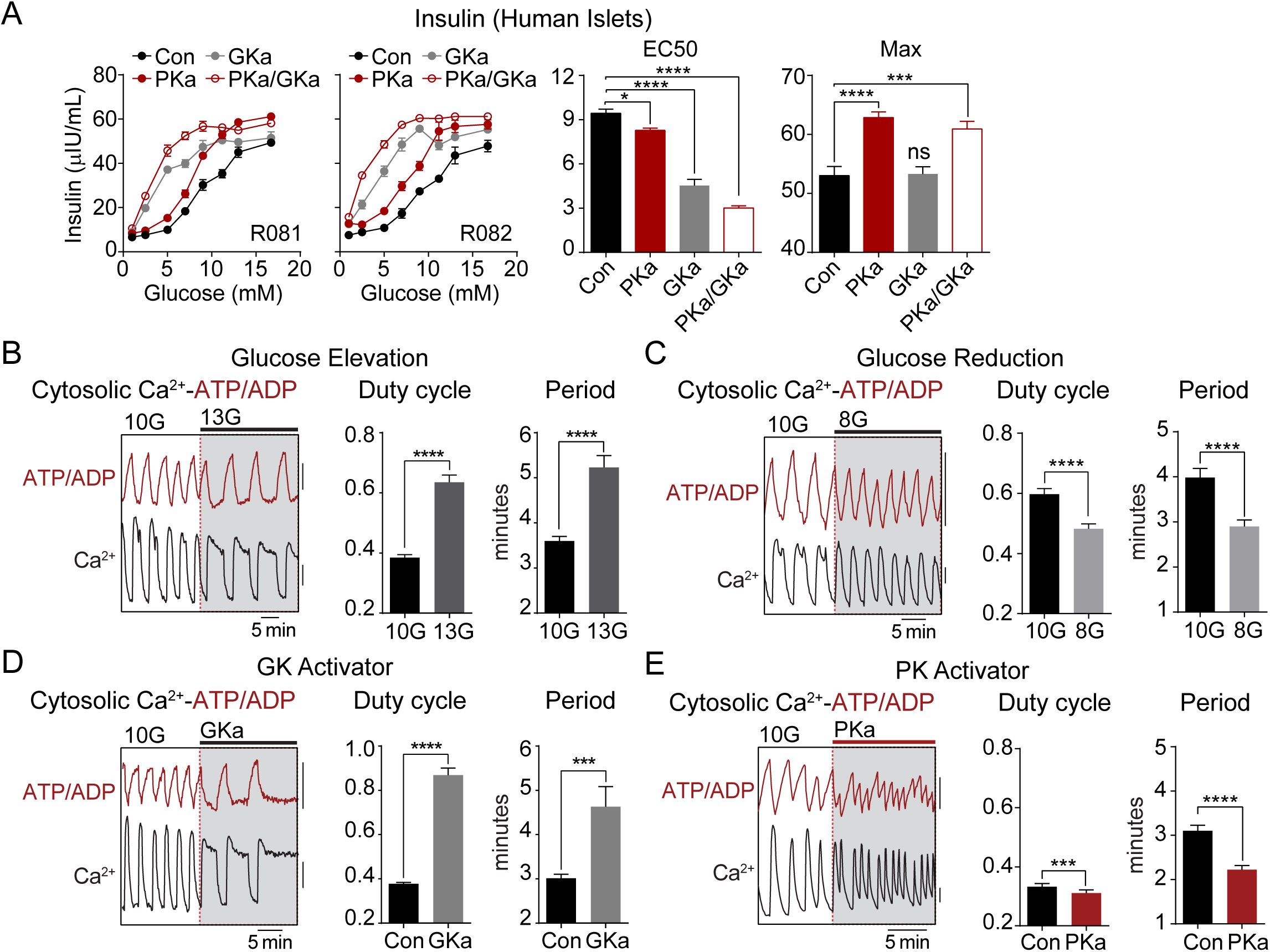
PK amplifies insulin secretion by a distinct mechanism from glucokinase, which controls the β-cell triggering pathway. (A) GKa and PKa potentiate insulin secretion in human islets (n = 8) in an additive manner. (B-E) Representative recordings and quantification of cytosolic calcium and ATP/ADP oscillations stimulated by 10 mM glucose (10G). (B) Duty cycle and period were increased by glucose elevation (n = 51). (C) Reducing the glucose concentration to 8 mM glucose (8G) reduced duty cycle and period (n = 53). (D) 500 nM GK activator (GKa) RO-0281675 increased duty cycle and period (n = 37). (E) 10 µM PK activator TEPP-46 did not change duty cycle but decreased period (n = 62 islets from 4 mice). FuraRed (Ca^2+^), black scale bar = 0.01 (B, D) or 0.1 (C, E) R430/500; Perceval-HR (ATP/ADP), red scale bar = 0.01 R500/430. Data are shown as mean ± SEM. *p < 0.05, ***p < 0.001, ****p < 0.0001 by 1-way ANOVA (A) or paired t-test (B-E).

To evaluate the β-cell triggering pathway, we performed simultaneous measurements of cytosolic ATP/ADP and calcium oscillations in mouse islets. As the primary glucose-dependent parameter, the calcium duty cycle (fractional time in the active phase of an oscillation) reflects processes that act via K_ATP_ (Henquin, 2009). Raising glucose from 10 to 13 mM increased the calcium duty cycle (**Figure 3B**), while lowering glucose from 10 to 8 mM had the opposite effect (**Figure 3C**). GK activator (0.5 μM RO-0281675) (Grimsby et al., 2003) increased the duty cycle in a similar fashion to raising glucose (**Figure 3D**). Failing to match the effects of glucose or GK activator, PKa actually lowered the calcium duty cycle (**Figure 3E**) despite increasing insulin secretion (**Figure 2**). At the same time, PKa accelerated ATP/ADP cycling and reduced the time between adjacent depolarizations, as indicated by a marked, 28% reduction in the calcium oscillation period (**Figure 3E**). Thus, while PK is incapable of matching the effects of GK on glucose uptake and calcium duty cycle, we observed two GK-independent effects: PK modulates the frequency of membrane depolarization and controls the amplitude of insulin secretion.

### The GK-independent actions of PK are powered by mitochondrial anaplerosis

An important observation is that PK activators did not stimulate at glucose concentrations < 5mM (**Figures 2I and 3A**). Even as PK is sufficient to close K_ATP_ channels (**Figure 1**), the insulin secretion and calcium imaging data confirm that when glucose is the substrate then PK is dependent on glycolytic PEP that is under the metabolic control of GK (**Figure 3**). But importantly, PEP can arise from *both* the glycolytic enolase reaction as well as from anaplerosis contributing to PEPCK-M mediated cataplerosis known as the PEP cycle (**Figure 4A**) (Jesinkey et al., 2019; Stark et al., 2009). To understand the mechanism by which PEP cycling interacts with PK, we utilized substrates that supply PK with PEP independently of glycolysis. Succinate, or the membrane-permeant succinic acid methyl ester (SAME), is a pure anaplerotic substrate (i.e. it is not a direct source of acetyl CoA) that can expand the oxaloacetate pool to feed PEPCK-M (Stark et al., 2009). In human islets, SAME required sub-stimulatory glucose to stimulate insulin secretion but shifted the glucose sensing curve to the left in a similar manner to GK activators. SAME also displayed an amplifying behavior relative to controls and increased maximal secretion to a similar degree as PK activation (**Figure 4B**). The combination of SAME and PK activation were also additive, but predominantly increased maximal secretion. For SAME to be responsive to PKa, it must enhance the supply of PEP. This fits with the observation, shown in the companion paper, that SAME requires PEPCK-M to increase calcium and insulin secretion (Abulizi et al., 2019) and confirms in human islets that, like glycolysis, mitochondrial anaplerosis-cataplerosis can provide a source of PEP to PK that, in turn, is responsive to PK activation.

**Figure 4.**
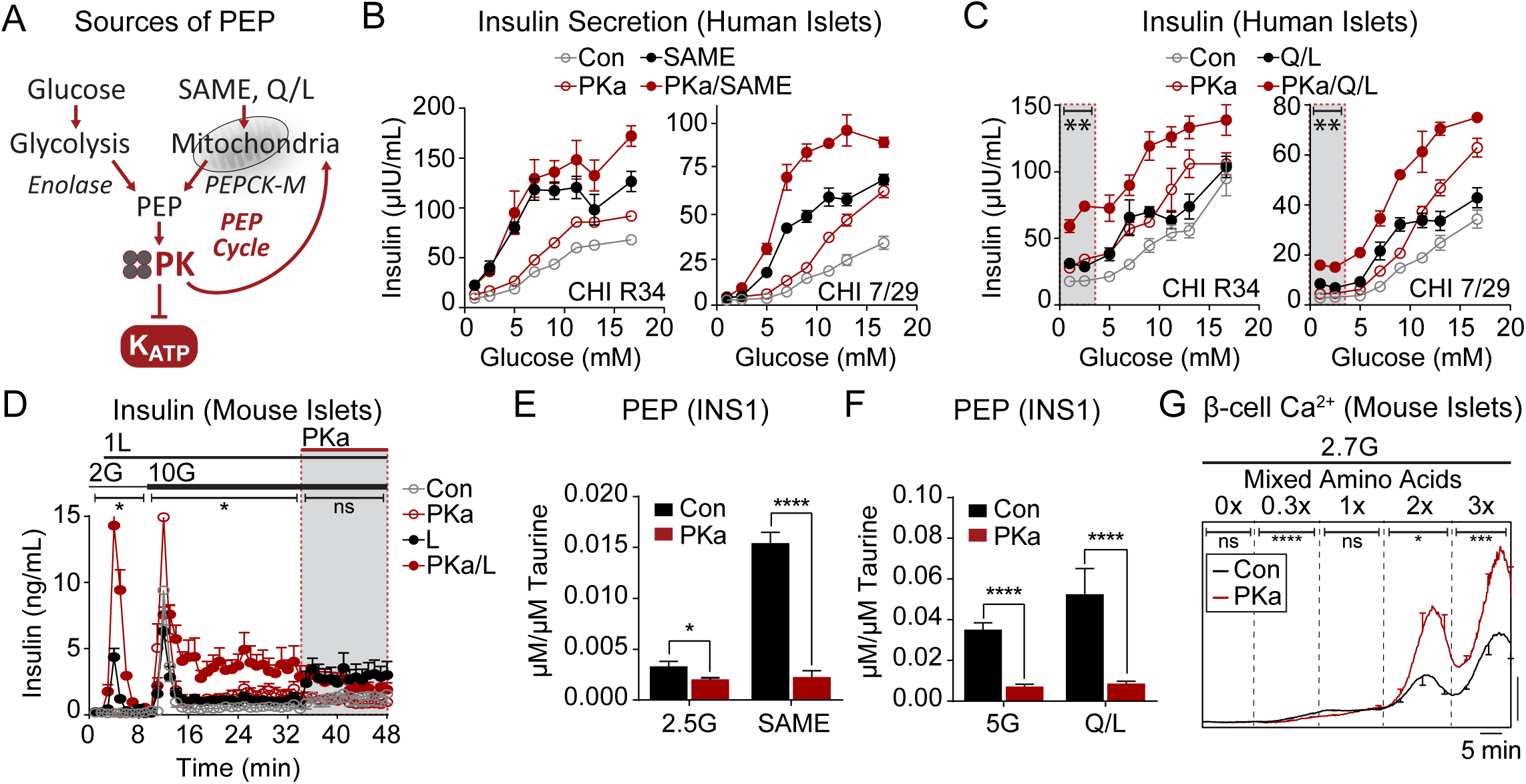
The GK-independent action of PK is powered by mitochondrial anaplerosis. (A) Cartoon depicting the sources of phosphoenolpyruvate (PEP) from glycolysis and mitochondrial anaplerosis. SAME (monomethyl succinate), PEPCK-M (mitochondrial phosphoenolpyruvate carboxykinase). (B) Insulin secretion from 2 human islet donors with glucose (1-16.7 mM), 10 mM monomethyl succinate (SAME), and 10 µM PKa as indicated. (C) Insulin secretion from 2 human islet donors with glucose (1-16.7 mM), 4 mM glutamine plus 10 mM leucine (Q/L), and 10 µM PKa as indicated. (D) Insulin release from mouse islets in the presence of 2 and 10 mM glucose (2G, 10G), 1 mM leucine (1L), and 10 µM PKa as indicated (n = 8 mice). (E) Concentration of PEP in INS1 832/13 cells (n = 6) in response to 2.5 mM glucose (2.5G) and 10 mM monomethyl succinate (SAME) in the absence or presence of 10 µM PKa. (F) Concentration of PEP in INS1 832/13 cells (n = 6) in response to 5 mM glucose (5G) and 4 mM glutamine plus 10 mM leucine (Q/L) in the absence or presence of 10 µM PKa. (G) Representative average β-cell calcium in the absence or presence of PKa and in response to an amino acid ramp at 2.7 mM glucose (2.7G) in mouse islets. PKa increased β-cell calcium in 2.7G (Con, n = 19; PKa, n = 17). Data are shown as mean ± SEM. *p < 0.05, **p < 0.01, ***p < 0.001, ****p < 0.0001 by t-test. See also Figure S4 and Table S2.

At sub-stimulatory glucose concentrations, SAME ± PKa did not stimulate insulin secretion, suggesting a minimal requirement for an oxidizable carbon source such as AcCoA (**Figure 4B**). While glutamine by itself doesn’t stimulate insulin secretion, the combination of glutamine plus leucine (Q/L) provides a direct source of acetyl CoA (through the oxidation of leucine) and a source of anaplerotic α-ketoglutarate (via allosteric activation of glutamate dehydrogenase by leucine), subverting the need for glycolysis. Like SAME, Q/L shifted the glucose sensing curve to the left and the combination of Q/L and PKa amplified and maximized insulin secretion (**Figure 4C**). In contrast to SAME, Q/L could trigger insulin secretion at sub-stimulatory glucose concentrations that could be amplified by PKa (**Figure 4C, *inset box***). Interestingly, the augmentation of insulin secretion by Q/L shifted insulin secretion upward and was relatively constant across the whole glucose range, suggesting a fixed AcCoA addition by leucine. Consistently, in mouse islets, the synergistic effects of leucine and PKa on insulin secretion were clearly evident at basal and stimulatory glucose concentrations (**Figure 4D**). Importantly, these data suggest that PK activation does not require GK as long as an ample supply of both oxidizable and anaplerotic carbon are present.

At low glucose concentrations, when PEP cycling is minimal, the steady state PEP concentration is determined simply by its glycolytic supply and PK mediated clearance. Here, the activity of PKa on its target PK was confirmed by a reduction in the steady state PEP concentration (**Figures 4E and 4F**). The increase in PEP following the addition of SAME or Q/L indicates a PEPCK-M mediated influx into the PEP pool. This cataplerotic pool was drained by accelerating PEP clearance by PK activation.

We also determined if insulin secretion from anaplerotic metabolism was mediated by cytosolic calcium in a PK dependent manner. At low glucose, increasing the mixed amino acid concentrations progressively increased the mean calcium response in mouse islet β-cells, which reached a substantially higher level in the presence of PKa (**Figure 4G and Table S2**). The effect of PKa is progressively diminished as glucose concentrations are raised and glycolytic PEP is generated (**Figure S4**), suggesting redundancy in the triggering system. These data demonstrate that, like with insulin secretion (**Figures 4B, 4C**), mitochondrial PEP works through PK to increase calcium.

### PK activation does not alter steady-state metabolism

If interpreted through the lens of the consensus model, the observation that PK activation significantly increased insulin secretion is expected to be accompanied by a global increase in mitochondrial metabolism. It was surprising, then, that PK activation did not significantly increase oxygen consumption rates (OCR) in either INS1 832/13 cells or mouse islets (**Figures 5A and 5B**). Similarly, at 9 mM glucose, PK activation did not change the concentration of central carbon metabolites (**Figure 5C**) including PEP. The lack of a PKa-driven PEP reduction suggests that at higher glucose, glycolysis and PEP cycling are able to keep pace with PK. Since metabolite concentrations do not directly correlate with metabolic flux, a precision metabolic flux platform, MIMOSA (Mass Isotopomeric MultiOrdinate Spectral Analysis), which has a sensitivity to the impact on metabolism of as little as 0.5 mM glucose (Alves et al., 2015), was used to directly examine the mitochondrial pathways dependent on pyruvate. In an apparent paradox, despite increasing insulin secretion, independent measures using [U-^13^C_6_]-D-glucose for kinetic modeling (*v*) or steady state (Φ) calculations, PK did not significantly impact oxidative, PC-dependent anaplerotic, or PEPCK-M-dependent cataplerotic metabolic rates (**Figures 5D, 5E)**.

**Figure 5.**
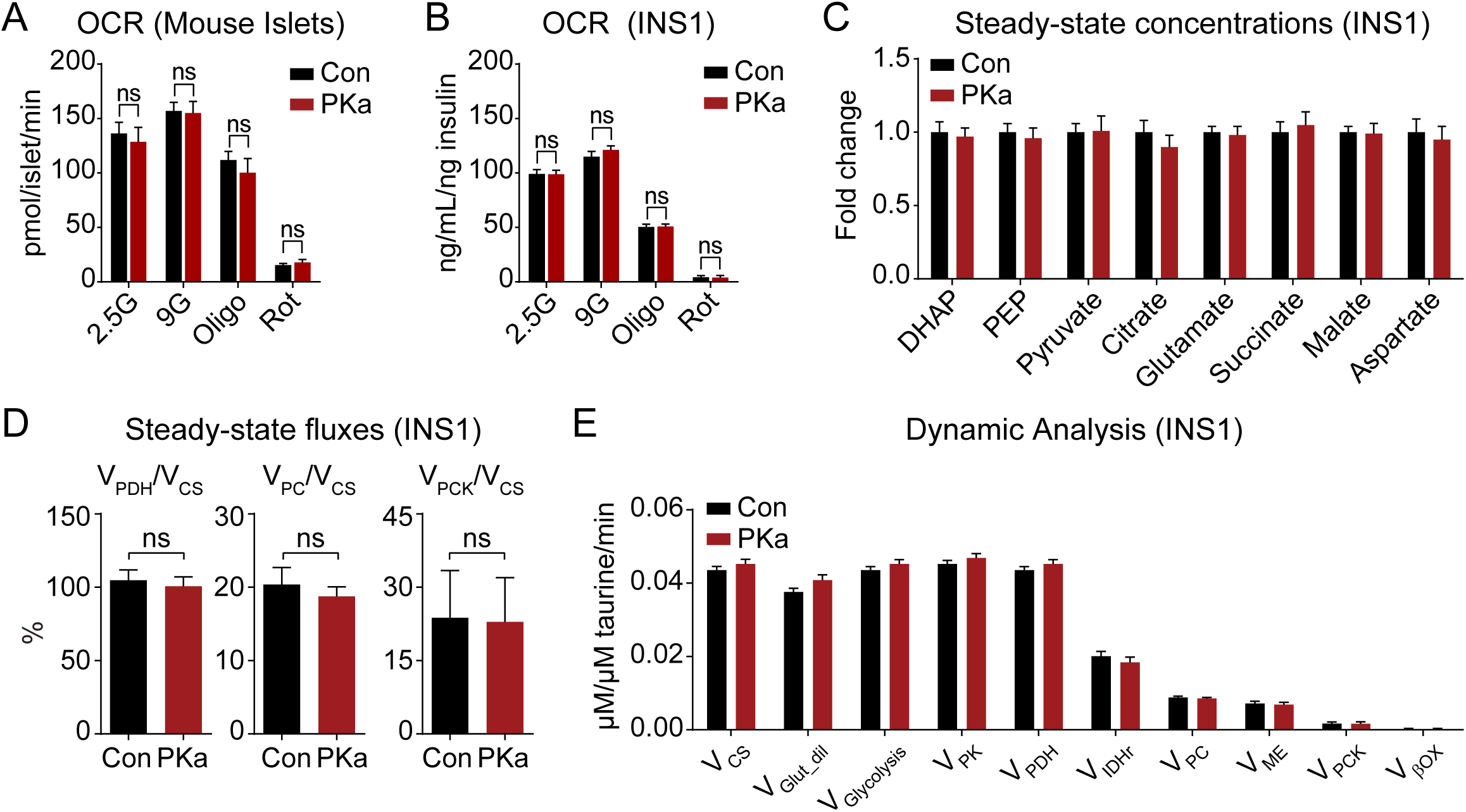
PK activation is independent of oxidative mitochondrial metabolism. (A) Oxygen consumption rate of mouse islets (n = 3) treated with 10 µM PK activator TEPP-46 (PKa, n = 5) or vehicle (Con, n = 4) and the acute addition of electron transport chain inhibitors (Oligo, 1 µM oligomycin; Rot, 1 µM rotenone). (B) Oxygen consumption rate of INS1 832/13 cells (n = 15) treated with 10 µM PK activator TEPP-46 (PKa) or vehicle (Con) and the acute addition of electron transport chain inhibitors (Oligo, 1 µM oligomycin; Rot, 1 µM rotenone). (C-E) Impact of PK activation on INS1 832/13 metabolic fluxes. (C) Steady-state concentrations of glycolytic and TCA cycle intermediates. (D) Fractional flux through pyruvate dehydrogenase (V_PDH_), pyruvate carboxylase (V_PC_), and phosphoenolpyruvate carboxykinase (V_PCK_) obtained from the steady-state analysis of enrichments following incubation with [U-^13^C_6_]glucose. (E) Absolute fluxes calculated by the mathematical analysis of the time-dependent accumulation of ^13^C-label into the glycolytic and mitochondrial intermediates (n = 6). Data are shown as mean ± SEM. Statistics calculated by t-test.

### By cyclically depriving mitochondria of ADP, PK activation restricts OxPhos

The lack of evidence in change in steady-state metabolism despite evidence of increased PK activity and insulin secretion suggests that the linear metabolic model that to this point had been assumed is too simplistic. If PKa changed the frequency of metabolic oscillations, then the flux measurements above (**Figure 5D, 5E**) would not be sensitive to the changes at steady state. Although PK is able to increase insulin secretion without changing the *time-averaged* oxidative fluxes, we did observe changes in *time-resolved* mitochondrial metabolism at a single islet level. PKa effectively reduced the cycling period of NAD(P)H oscillations, which primarily reflect mitochondrial NADH (Patterson et al., 2000), as well as mitochondrial membrane potential (ΔΨm) oscillations (**Figures 6B**), matching the effects of PKa on ATP/ADP and calcium cycling (**Figure 3E**). We also observed that PKa increased mitochondrial NAD(P)H fluorescence, and strongly increased ΔΨm, in the presence of both 9 and 2 mM glucose (**Figures 6B, 6C**). In particular, PK-induced ΔΨm hyperpolarization would be consistent with decreased OxPhos as would occur in “State 4-like” metabolism when ADP, rather than substrate, is limiting (**Figure 6A**).

**Figure 6.**
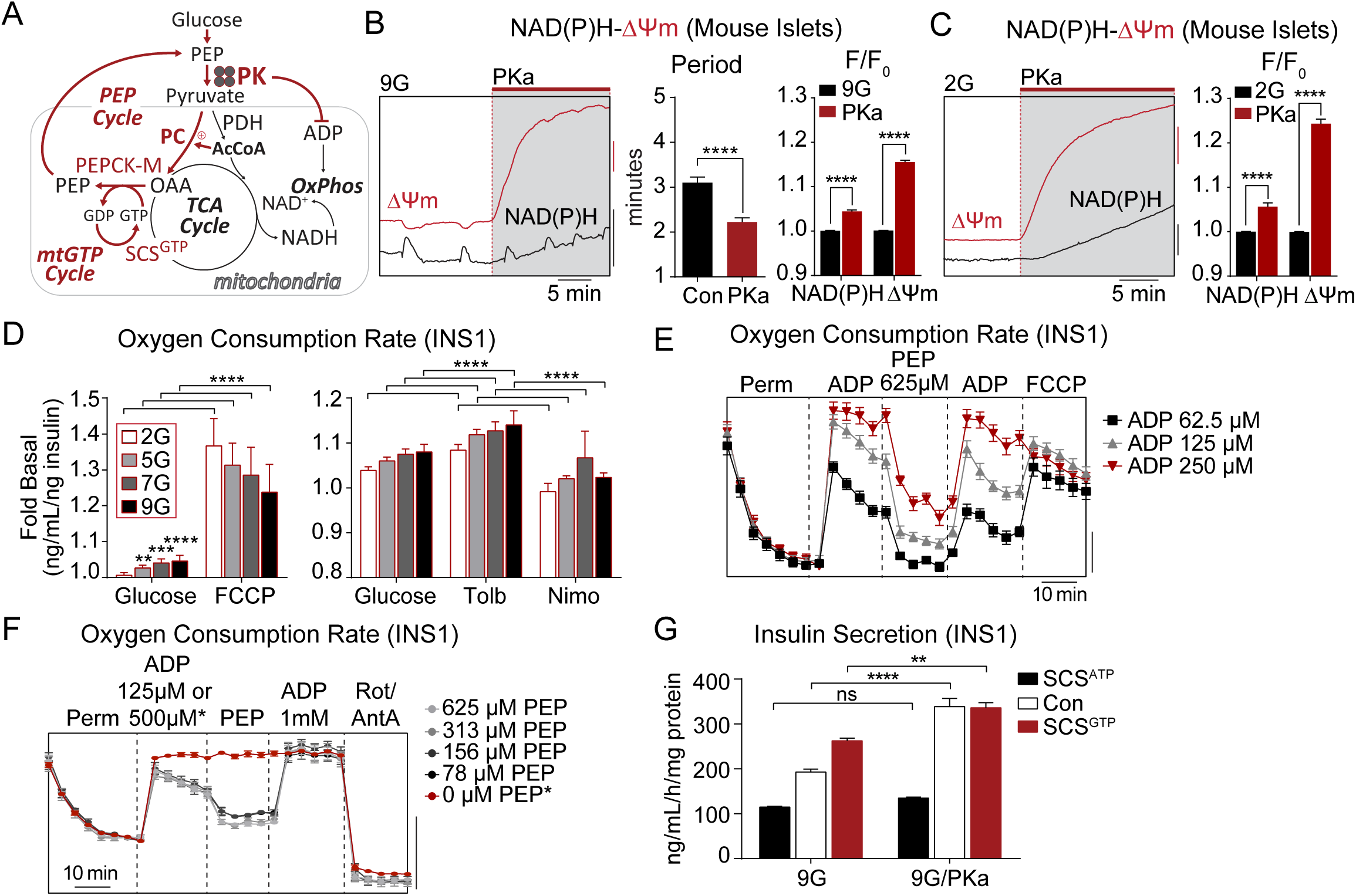
PK requires the mitochondrial PEP cycle to amplify insulin secretion. (A) Cartoon depicting the coordination between PK-mediated ADP lowering, inactivation of the electron transport chain (ETC), and increased mitochondrial GTP (mtGTP) and PEP cycling. PC (pyruvate carboxylase), PDH (pyruvate dehydrogenase), AcCoA (acetyl CoA), OAA (oxaloacetic acid), SCS^GTP^ (GTP-producing isoform of succinyl CoA synthetase), OxPhos (oxidative phosphorylation). (B) Representative recordings and quantification of NAD(P)H fluorescence and mitochondrial membrane potential (ΔΨm) oscillations stimulated by 9 mM glucose (9G) followed by acute application of PKa (Con, n = 36; PKa, n = 43). F/F_0_ indicates fluorescence normalized to baseline. (C) Representative recordings and quantification of NAD(P)H fluorescence and mitochondrial membrane potential (ΔΨm) stimulated by 2 mM glucose (2G) followed by acute application of PKa (Con, n = 21; PKa, n = 21). F/F_0_ indicates fluorescence normalized to baseline. (D) Oxygen consumption of INS1 832/13 cells (n = 6) treated with or without 10 µM FCCP, 100 µM tolbutamide, or 5 µM nimodipine at 2 mM, 5 mM, 7 mM, or 9 mM glucose. (E) Oxygen consumption of INS1 832/13 cells (n = 6) treated with Plasma Membrane Permeabilizer (Perm), ADP (62.5 µM, 125 µM, or 250 µM), 625 µM PEP, and 4 µM FCCP. Oxygen consumption rate, scale bar = 50 pmol/min. (F) Oxygen consumption of INS1 832/13 cells (n = 6) treated with Plasma Membrane Permeabilizer (Perm), ADP (125 µM or 500 µM, for 0 µM PEP condition), PEP (625 µM, 313 µM, 156 µM, 78 µM, or 0 µM) and Antimycin A and Rotenone (A/R, 10/5 µM). Oxygen consumption rate, scale bar = 1 pmol/min. (G) Insulin secretion from PKa- or vehicle-treated control INS1 832/13 cells (Con, n = 12), or INS1 832/13 cells stably overexpressing the ATP and GTP-producing isoforms of succinyl CoA synthetase (SCS^ATP^ and SCS^GTP^, respectively) (n = 6) in response to 9 mM glucose (9G). Data are shown as mean ± SEM. *p < 0.05, **p < 0.01, ***p < 0.001, ****p < 0.0001 by t-test (B-C), 2-way ANOVA (D), or 1-way ANOVA (G).

If β-cell mitochondrial respiration were purely substrate-limited, as predicted by the canonical model, then uncoupling should not increase respiration. Instead, ADP limitation was exposed when FCCP uncoupled the proton motive force from ATP synthase, without correlation to the ambient glucose concentration (**Figure 6D**). Consistently, increasing ATP hydrolysis (with tolbutamide) increased respiration, while reducing workload (with the further addition of nimodipine) had the opposite effect (**Figure 6D**). To directly test whether PK itself is capable of limiting OxPhos by ADP privation *in situ*, respirometry experiments were conducted in permeabilized INS1 832/13 cells in the presence of ADP and succinate (**Figure 6E**). While ADP dose-dependently stimulated State 3 respiration as would be expected, the further addition of PEP immediately reduced respiration consistent with a shift towards State 4. This inhibition was overcome by replenishment of ADP which increased respiration to its maximal uncoupled rate (**Figure 6E**). Considering the observed ADP privation for local control of K_ATP_ closure (**Figure 1**), and the saturable effect of PKa on ΔΨm at low glucose (**Figure 6B**), we considered that PK may also act locally to restrict OxPhos. Confirming this hypothesis, PEP instantaneously restricted respiration when present at sub-stoichiometric levels relative to ADP (**Figure 6F**). These data demonstrate the existence of a novel, functionally-defined metabolic compartment in which PK locally controls OxPhos through cytosolic ADP privation.

### PK requires the mitochondrial PEP cycle to amplify insulin secretion

The ability of PKa to intermittently restrict ΔΨm and NADH without any net impact on oxidative mitochondrial metabolism (**Figures 5, 6 and S4**) suggests a mechanism by which PK is able to transiently restrict PDH, and autocatalytically reinforce the anaplerotic fluxes dependent on PC and the cataplerotic PEPCK-M reactions that refuel the cytosolic PEP pool (Alves et al., 2015; Kibbey et al., 2007) (**Figure 6G**). PEPCK-M is dependent on mitochondrial GTP (mtGTP) produced from the GTP-specific isoform of the TCA cycle enzyme succinyl CoA synthetase (SCS). Both the GTP isoform (SCS-GTP) and the ATP isoform (SCS-ATP) are catalytically dependent on a shared alpha subunit, so silencing or overexpressing one or the other isoform can modulate mtGTP synthesis and therefore PEPCK-M activity (Jesinkey et al., 2019; Kibbey et al., 2007). To determine whether PKa requires the PEP cycle to stimulate insulin secretion, we took advantage of INS1 832/13 cells stably overexpressing SCS-GTP or SCS-ATP (Jesinkey et al., 2019). By comparison to the parental INS1 832/13 cells, GSIS was enhanced in SCS-GTP cells and blunted in SCS-ATP cells (**Figure 6G**). In the SCS-ATP cells, the effect of PKa to enhance insulin secretion was completely blocked, by comparison to control cells where secretion was increased by 75%. In SCS-GTP cells, PKa only modestly increased secretion, in this case because insulin secretion was already near the PKa-stimulated maximum (**Figure 6G**). These results identify an essential role for the PEP cycle in mediating the effect of PK on insulin secretion.

### Evidence for a 2-state model of oscillatory β-cell metabolism

Using molecular biosensors, we characterized the real-time phasic relationship between PK activity, mitochondrial and plasma membrane potentials, K_ATP_ conductance, cytosolic calcium, lactate and glutamate, mitochondrial NAD(P)H, and the cytosolic ATP/ADP ratio (**Figures 7A-G**). In parallel, we used time-resolved stable isotopomer enrichment analysis to measure the oscillatory fluxes of PC relative to PDH (**Figure 7H**). Based on these measurements, we propose a 2-state model of β-cell metabolism (**Figure 7I**) describing the relationship of PK activation to β-cell metabolic and electrical oscillations.

**Figure 7.**
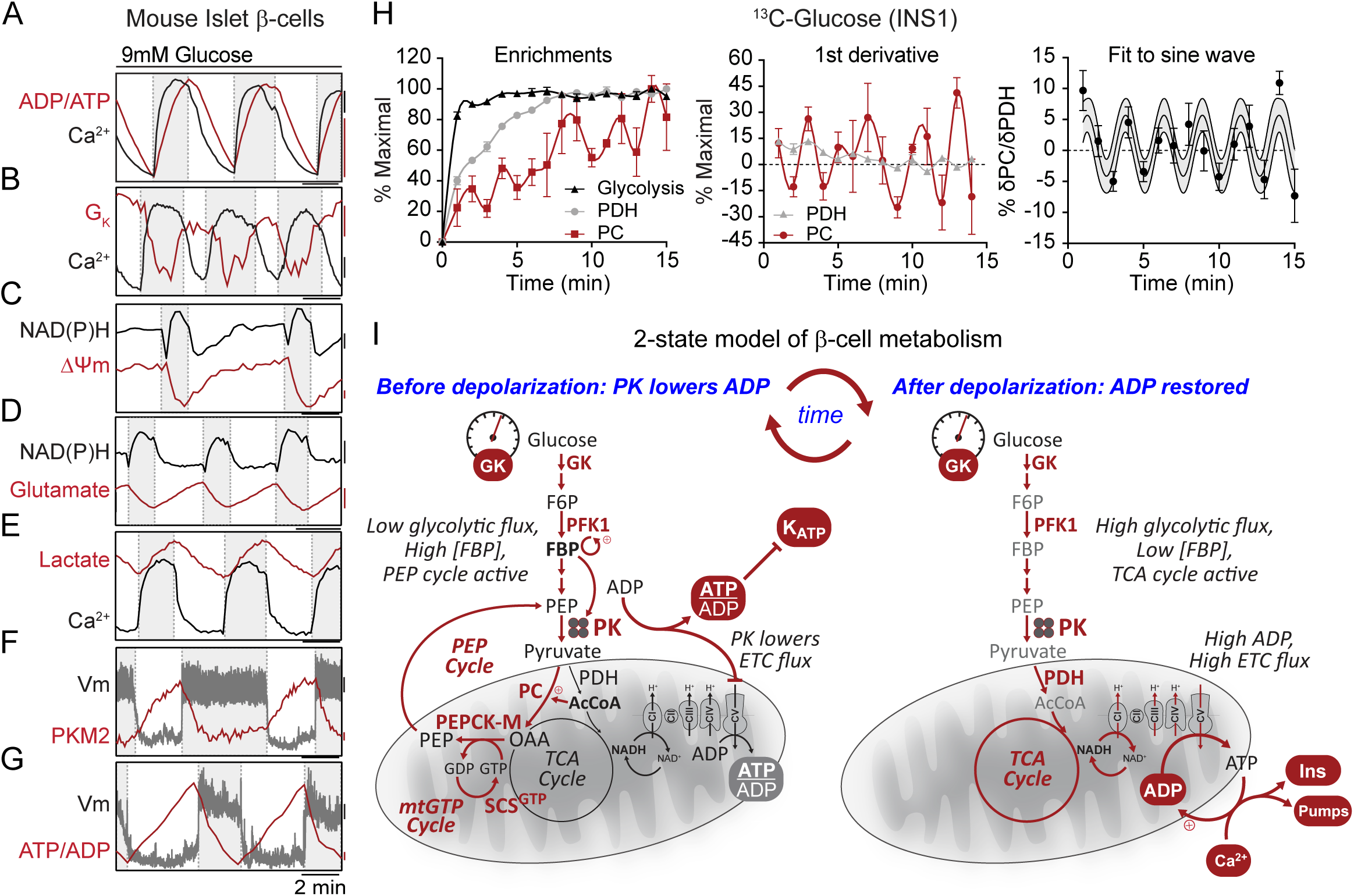
Evidence for a 2-state model of oscillatory β-cell metabolism. (A-G) Metabolic and electrical oscillations in islet β-cells stimulated with 10 mM glucose. ATP/ADP, calcium, glutamate, lactate, and PKM2 are cytosolic parameters. NAD(P)H and ΔΨm are mitochondrial parameters. Plasma membrane potential (Vm) and potassium conductance (G_K_) are electrical parameters. Silent phase, white boxes; active phase, shaded boxes. (H) Antiphase oscillations in flux through PDH and PC in INS1 832/13 cells following the addition of 9 mM glucose. (I) Model of oscillatory β-cell metabolism with 2 states (triggering vs. secretory) separated by membrane depolarization. Before depolarization, PK lowers ADP, reducing flux through the ETC (ADP-starved, “State 4-like”) and the TCA cycle while activating the PEP cycle. After PK lowers ADP sufficiently to close K_ATP_ channels, workload in the form of ATP hydrolysis restores ADP (e.g. by exocytosis and pumps), increasing flux through the ETC (ADP replete “State 3-like”), the TCA cycle, and glycolysis. Note that recruitable PK isoforms (M2 and L) are active before depolarization, when glycolytic flux is low and the FBP concentration is high.

β-cells have electrically silent and active phases (white versus grey boxes in **Figures 7A-G**) characterized by the square waves of the plasma membrane potential (V_m_). The beginning and ending of each oscillatory phase coincide with the peaks and troughs of ATP/ADP. The metabolic and electrical features described below start with the 1) silent phase with 2) progressive transition into State 4-like respiration with activation of the anaplerotic/cataplerotic PEP cycle 3) to trigger depolarization and 4) activate OxPhos:

#### 1) The electrically silent phase

The transition into the electrically silent phase begins when ADP levels rise sufficiently to re-open K_ATP_ channels and hyperpolarize the plasma membrane as well as close voltage-dependent calcium channels (**Figures 7A, B, F, G**). The sudden decrease in metabolic demand reduces ATP hydrolysis rates and allows glycolysis and OxPhos to clear calcium and raise ATP/ADP (**Figures 7A, B, G**).

#### 2) State-4 like respiration

The progressive lowering of ADP by both OxPhos and glycolysis eventually slows the TCA cycle. This is identified by the asymptote of ΔΨ_m_ and mitochondrial NAD(P)H midway through the silent phase that hallmarks State 4 respiration (**Figure 7C**), as previously observed by Kennedy and colleagues who measured oxygen (Kennedy et al., 2002). As the TCA cycle stalls, mitochondrial metabolites accumulate so that AcCoA can activate the PC-dependent generation of OAA observed by stable isotope patterning (**Figure 7H**). Through the mass action of OAA synthesis and high mitochondrial NADH, the glutamate-aspartate shuttle slows (as indicated by cytosolic glutamate accumulation) (**Figure 7D**). The sn-glycerol-3-phosphate shuttle would also be expected to slow in the face of a higher ΔΨ_m_. In the presence of continuous GK activity but with GAPDH slowing (from increased cytosolic NADH/NAD^+^), the levels of FBP increase to activate PK allosterically (**Figure 7F**). With GAPDH slowed, PEP cycling via PEPCK-M would continue to lower ADP. The lowering of lactate can be explained in the setting of slowed pyruvate dehydrogenation and reduced glycolysis (**Figure 7E**). A high mitochondrial ATP/ADP ratio reverses SCS-ATP to provide additional succinyl CoA for enhanced SCS-GTP-mediated mtGTP synthesis. PEPCK-M utilization of OAA and mtGTP supports rapid mitochondrial PEP synthesis to further lower ADP in the cytosol until K_ATP_ channels close, marking the end of this phase **(Figures 7A, B, F, G)**. In short, PK-mediated ADP lowering slows OxPhos to divert mitochondrial pyruvate away from oxidation and into PEP synthesis. Such feed forward metabolism continues to accelerate PEP cycling until ATP/ADP elevation triggers K_ATP_ closure.

#### 3) The electrically active phase

*After* membrane depolarization and Ca^2+^ influx, during the electrically active phase of an oscillation (the shaded boxes in **Figures 7A-G**), insulin exocytosis occurs. This active phase persists as long as ADP is maintained below the K_ATP_ threshold.

#### 4) State 3-like respiration is sustained by the ETC

During this active phase, the sudden surge in metabolic workload (e.g., by Ca^2+^ pumping and exocytosis, **Figure 6D**) accelerates ATP hydrolysis (**Figure 7A**). The combination of a high ADP load and a spike in mitochondrial calcium activates OxPhos (e.g., ATP synthase, TCA cycle dehydrogenases, etc.) to restore “State 3-like” OxPhos. Active OxPhos then consumes (depolarizes) ΔΨm and mitochondrial NADH (**Figure 7C**), and an active TCA cycle lowers AcCoA, transitioning mitochondria from anaplerotic PC to favor oxidative PDH (**Figure 7H**). Concurrently, the glutamate-aspartate shuttle reactivates with cytosolic glutamate decreasing in the face of a more receptive mitochondrial NADH/NAD^+^ ratio and diminished PC anaplerosis (**Figure 7D**), and elevated glycolytic flux lowers the cytosolic NADH/NAD^+^ ratio as is reflected by the increase in lactate (**Figure 7E**). As glycolytic intermediates are lowered, the drop in FBP inactivates PKM2 and PKL (**Figure 7F**). This highly oxidative state, in which calcium and OxPhos synergize, most resembles the β-cell consensus model where the capacity to support the ATP/ADP ratio sustains depolarization in the face of high ATP demand. This electrically active phase ends as ATP hydrolysis exceeds the capacity of mitochondria to keep ADP low and K_ATP_ channels re-open and the plasma membrane repolarizes.

As long as glucose remains elevated, oscillations between these two states persist.

## DISCUSSION

These data provide evidence that PEP hydrolysis by PK, rather than OxPhos, provides the energetic push required to raise ATP/ADP beyond the threshold for K_ATP_ channel closure. The functional linkage between PK and K_ATP_ in β-cells is consistent with prior studies in cardiac myocytes, where PK closes K_ATP_ channels (Weiss and Lamp, 1987; Dhar-Chowdhury et al., 2005). Importantly, PEP can close K_ATP_ channels in isolated membrane patches despite the presence of the strong K_ATP_ channel-opener ADP, indicating that PK can influence the metabolic microenvironment near the channel. The demonstration that PK is able to restrict OxPhos by locally depriving the mitochondria of ADP provides a second example of β-cell metabolic compartmentation in which ADP is the signal carrier, and solidifies the importance of glycolytic ATP/ADP regulation.

Our data do not undermine the previously described metabolic control exerted by GK that is limited to glycolysis (Matschinsky and Ellerman, 1968). Real-time imaging of calcium oscillations reaffirmed that the threshold for membrane depolarization is set by GK and observable in the duty cycle (Henquin, 2009), and identify a second mode of triggering based on mitochondrial PEP biosynthesis. Based on the canonical OxPhos-only model, PK activation should not be able to increase GSIS since GK activity determines the glycolytic and, consequently, mitochondrial flux rates. It is important to note that anaplerosis has been identified as an important component of metabolic coupling without a clear mechanism (Prentki et al., 2013). Islet perifusion studies revealed that, independently of initiating membrane depolarization (*i*.*e*. in the presence of KCl), the dynamically regulated, allosterically recruitable PK appears to play a dominant role in amplifying secretion. This is reinforced by the requirement for an AcCoA source for triggering but an anaplerotic carbon source for metabolic amplification in human and mouse islets. Thus, a key finding here is that PK underlies a separate mode of β-cell glucose-sensing downstream of GK that may provide a mechanism dependent upon anaplerosis. While targeting OxPhos in β-cells has not been successful therapeutically, preclinical data suggest the anaplerotic mechanism may be of potential benefit (Abulizi et al., 2019).

Surprisingly, PK activation did not significantly increase time averaged mitochondrial metabolism. Instead, it increased the frequency rather than the duration of the calcium pulses. Human islet secretion measurements likewise indicated that PK acts as a fuel-dependent amplifier that does not significantly lower the threshold for insulin secretion. The most straightforward explanation for metabolic amplification is that PK is limiting during periods of high glycolytic flux, such as the state induced by KCl and high calcium, when FBP levels are low and the recruitable PK isoforms are least active. PK activators stabilize the active enzyme and circumvent the need for FBP. These data from intact cells corroborate prior studies showing that ADP and PEP (but not pyruvate) are sufficient for biphasic insulin secretion in membrane-permeabilized islets, including in the presence of rotenone (Pizarro-Delgado et al., 2016).

Incorporating the data herein, we propose a 2-state model of β-cell metabolism describing the relationship of PK to the intrinsic β-cell metabolic oscillations. In this model, ADP availability switches between a state where anaplerosis is high and one where OxPhos is high. Here, the importance of metabolic control exerted by ADP stems from the simple fact that ADP must be reduced to low μM concentrations for K_ATP_ channels to close (Tarasov et al., 2006) and that mitochondrial ATP synthase cannot run without ADP (Chance and Williams, 1955). Transition between the 2 states is toggled by the large amount of ATP hydrolysis associated with membrane depolarization, pumps, and vesicle fusion (Nicholls, 2016; Affourtit et al., 2018). A key advantage of allosteric PK recruitment is the ability to reinforce metabolic oscillations in response to metabolic regulators like FBP (Merrins et al., 2013, 2016). Consequently, the apex of PK activity occurs just prior to membrane depolarization, at the nadir of OxPhos, which resumes following depolarization-initiated ATP hydrolysis. This back and forth between an electrically silent triggering phase and an active oxidative secretory phase allow β-cell mitochondria to move between PEP biosynthesis and OxPhos. The mitochondrial contribution of PK substrate from the PEP cycle during the silent phase boosts PEP production beyond what is achievable by glycolysis alone to enhance the cytosolic ATP/ADP-generating capacity of PK in the triggering phase.

Paradoxically, PK recruitment occurs in parallel with the slowing of glycolytic flux. However, this recruitment occurs in the setting of continuous GK flux, when FBP levels rise and flux through PC increases, rerouting pyruvate flux through PEPCK-M (Jesinkey et al., 2019; Kibbey et al., 2007; Stark et al., 2009). Prior estimations of PEP cycling showed rates could reach ∼40% of the total PK flux in islets (Stark et al., 2009). These estimations did not consider that oscillatory metabolism divides OxPhos and anaplerosis into separate phases. As such, mitochondrial PEP cycling could potentially explain nearly all of PK activity during the electrically silent phase.

The 2-state model recognizes that the PEP cycle primarily triggers via PK while OxPhos primarily sustains the active phase. As both processes generate ATP, their individual contributions may not be fully separable in either the time or compartment domains, and both processes are necessary to achieve maximal secretion. However, the improper timing of OxPhos relative to membrane depolarization (**Figure 7** and (Jung et al., 2000)) raises the question of whether it is bioenergetically feasible that mitochondria, even if strategically positioned at the plasma membrane, could lower ADP sufficiently to close K_ATP_ channels, as PK was demonstrated to do. Our data are supported by observations of high ATP generation at the plasma membrane relative to the cytosol and mitochondria (Kennedy et al., 1999). It remains to be determined if there is a point at saturating glucose levels where, through either glycolysis and/or the PEP cycle, PK can locally deplete ADP sufficient to close K_ATP_ channels while other mitochondria at sites of active ATP hydrolysis have adequate ADP to conduct OxPhos simultaneously. This work only shows that it is possible for PEP cycle-supported, PK-mediated mitochondrial and K_ATP_ channel ADP privation to occur.

Although many qualitative β-cell bioenergetics studies have shown that ADP supply and demand are critical for OxPhos and insulin secretion (**Figure 6D** and (Ainscow and Rutter, 2002; Doliba et al., 2003; Panten et al., 1986; Sweet et al., 2004; Affourtit et al., 2018)), quantitative metabolic control analyses in β-cells have remained lacking (Affourtit et al., 2018; Nicholls, 2016). Since tools to quantify the phosphorylation potential and ATP flux currently rely on temporal averaging (Affourtit et al., 2018), it is not yet possible to calculate the relative contribution of PK and OxPhos to K_ATP_ closure. We also lack the means to determine how PK-driven ATP/ADP cycling bioenergetically supports increased exocytosis relative to OxPhos. Approaches will need to be developed that can absolutely quantify adenine nucleotides and their sources in a subcompartment-specific and time-resolved way. However, reevaluation of such β-cell bioenergetics may have similar application to other metabolic cycles and pathways that have been proposed (Schuit et al., 1997; Farfari et al., 2000; Joseph et al., 2006; Jensen et al., 2008; Prentki et al., 2013) but are not addressed in this simplified 2-state model.

This new understanding of the native role of PK in β-cell metabolism has broad implications in non-native environments where it has been coopted, such as in cancer (Dayton et al., 2016; Israelsen et al., 2013). It may be particularly advantageous for cancer or other dividing cells to shut down oxidative phosphorylation in order for the mitochondria to synthesize, rather than oxidize, building blocks. Time-resolved stable isotope measurements identify that OxPhos and anaplerosis are anti-phase with each other and suggests that mitochondria choose not to oxidize and synthesize simultaneously. In light of this, the phenomenon of PK-associated Warburg metabolism, while often considered for its ATP-generating capacity, may be even more important for its ADP lowering capacity (Harris and Fenton, 2019). Just as in β-cells, ADP deprivation may shift mitochondrial activity from OxPhos into the biosynthesis of essential nutrients needed to support cell division, or the generation of reactive oxygen species used for signaling. Of note, cells with more robust PEP cycling have an increased mass of elongated mitochondria while those deficient have fewer more fragmented mitochondria (Jesinkey 2019). The ability of PK isoforms to oscillate may also provide additional advantages not considered here, e.g. intermittently reducing membrane potential to reduce free radical damage that might occur if mitochondria were perpetually kept in a ‘State 4-like’ situation. In this context, activating PK may have some potential therapeutic benefits in certain situations where Warburg metabolism is observed.

Finally, the ability of pharmacologic PK activators to raise insulin secretion has broad conceptual implications for type 2 diabetes therapies. Not only is PK identified as a novel target for diabetes therapy, but we demonstrate that, for a given level of glucose, the secretory pathway can be internally remodeled to tune the glucose responsiveness of healthy and diseased human β-cells. Currently available drugs that increase insulin secretion by stimulating glucose uptake (e.g. GK activators) elevate the metabolic workload on each β-cell (Porat et al., 2011) and can lead to glucotoxic-like damage (Nakamura and Terauchi, 2015). Drugs that increase insulin secretion by directly triggering membrane depolarization such as sulfonylureas, while invaluable for treating some forms of MODY (Kim, 2015), decouple β-cell nutrient sensing with insulin secretion and increase the risk of hypoglycemia. PK-dependent remodeling of the β-cell metabolic pathways could lead to a treatment for diabetes that avoids these pitfalls.

## ACKNOWLEDGEMENTS

We thank Sam Stephens for reproducing our PK activator mouse GSIS studies. We thank Dawn Davis and Arnaldo de Souza for providing human islets. We thank David Zenisek, Bhupesh Mehta, Colin Nichols, Maria Remedi, Conor McClenaghan, and Christopher Emfinger for assistance with K_ATP_ measurements. The Merrins laboratory gratefully acknowledges support from the American Diabetes Association (1-16-IBS-212), the NIH/NIDDK (K01DK101683 and R01DK113103), the NIH/NIA (R21AG050135 and R01AG062328), the Wisconsin Partnership Program, and the Central Society for Clinical and Translational Research. HRV received a postdoctoral fellowship from the American Diabetes Association (1-17-PDF-155), HRF received a postdoctoral fellowship from HRSA (T32HP10010), and we acknowledge Dudley Lamming for contributing support for EP from R01AG062328. The Kibbey laboratory gratefully acknowledges NIH/NIDDK (R01DK092606, R01DK110181 and K08DK080142), CTSA (UL1RR-0024139), and DRC (P30DK045735). JEC is funded by the American Diabetes Association (1-18-JDF-017). MEC received a postdoctoral fellowship from NIH/NIDDK (F32DK116542). This work utilized facilities and resources from the William S. Middleton Memorial Veterans Hospital and does not represent the views of the Department of Veterans Affairs or the United States Government.

## AUTHOR CONTRIBUTIONS

MJM and RGK conceived the study and wrote the paper with SLL. SLL, RLC, and HRF performed the main body of experiments with assistance from TH, EP, CP, HRV, TCA, XZ, MEC, and IJ. MJM, RGK, CSN, JEC, and CJT provided resources. All authors interpreted the data and edited the manuscript.

## EXPERIMENTAL PROCEDURES

### Mouse Islet Isolation, Cloning, and Adenoviral Delivery of Biosensors

C57BL/6J mice were sacrificed via cervical dislocation and islet isolations were carried out as previously described (Gregg et al., 2016). Islets were cultured overnight in RPMI1640 supplemented with 10% (v/v) fetal bovine serum (Invitrogen A31605), 100 units/ml penicillin and 100 µg/ml streptomycin (Invitrogen). Genetically-encoded sensors for glutamate (Addgene) (GltI253-cpGFP.L1LV/L2NP, K_d_ = 107 μM) (Marvin et al., 2013) and lactate (Addgene) (San Martín et al., 2013) were cloned by Gibson Assembly (New England Biolabs) into a modified pENTR-DS shuttle vector (Invitrogen) containing the rat insulin promoter (RIP) and rabbit β-globin intron as in a previous study (Merrins et al., 2013). ClonaseII/Gateway was then used to prepare the full-length adenoviral construct in pAd/PL-DEST (Invitrogen). This procedure was used previously to generate adenovirus for β-cell specific Perceval-HR ATP/ADP biosensors (Merrins et al., 2016) and PKAR PKM2 biosensors (Merrins et al., 2013). For adenoviral delivery of the insulin promoter-driven biosensors, islets were infected immediately post-isolation with high-titer adenovirus for 2 hours at 37°C, then moved to fresh media overnight.

### Timelapse Imaging

For measurements of cytosolic Ca^2+^, islets were pre-incubated in 2.5 µM FuraRed (Molecular Probes F3020) in islet media for 45 min at 37°C before they were placed in a glass-bottomed imaging chamber (Warner Instruments) mounted on a Nikon Ti-Eclipse inverted microscope equipped with a 20X/0.75NA SuperFluor objective (Nikon Instruments). The chamber was perfused with a standard external solution containing 135 mM NaCl, 4.8 mM KCl, 2.5 mM CaCl_2_, 1.2 mM MgCl_2_, 20 mM HEPES (pH 7.35). The flow rate was 0.4 mL/min and temperature was maintained at 33°C using solution and chamber heaters (Warner Instruments). Excitation was provided by a SOLA SEII 365 (Lumencor) set to 10% output. Single DiR images utilized a Chroma Cy7 cube (710/75x, T760lpxr, 810/90m). Excitation (x) or emission (m) filters (ET type; Chroma Technology) were used in combination with an FF444/521/608-Di01 dichroic beamsplitter (Semrock) as follows: FuraRed, 430/20x and 500/20x, 630/70m (R430/500); NAD(P)H, 365/20x, 470/24m; Rhodamine-123 and Glutamate, 500/20x, 535/35m; and Perceval-HR, 430/20x and 500/20x, 535/35m (R500/430). Fluorescence emission was collected with a Hamamatsu ORCA-Flash4.0 V2 Digital CMOS camera every 6 s. A single region of interest was used to quantify the average response of each islet using Nikon Elements and custom MATLAB software (MathWorks).

### Electrophysiology

β-cell Ca^2+^ current and exocytosis were measured as in (Merrins and Stuenkel, 2008) with minor changes. Briefly, a Sutter MP-225 micromanipulator was used together with a HEKA EPC10 patch-clamp amplifier (Heka Instruments, Bellmore, NY) in the whole cell patch-clamp configuration to record Ca^2+^ current from intact islets perfused with standard external solution (above). Pipette tips were filled with an internal solution (in mM: 125 Cs-glutamate, 10 CsCl, 10 NaCl, 1 MgCl_2_·6H_2_O, 0.05 EGTA, 5 HEPES, 0.1 cAMP, 3 MgATP; pH 7.15 with CsOH) and 5 μM PKa or vehicle (0.1% DMSO) was added to this as indicated for dialysis into the cell. After membrane rupture, islet β-cells were identified by size (>5.5 pF), and after 1 min Ca^2+^ current was quantified from a 15 ms depolarization from −70 to 0 mV using a P/4 leak subtraction protocol. After 1 additional min, exocytosis was stimulated by activating VDCCs with a series of ten 500 ms membrane depolarizations from −70 to 0 mV. Capacitance responses (fF) and Ca^2+^ currents (pA) were normalized to initial cell size (pF). Islet KATP conductance was measured as described previously (Gregg et al., 2016). For single channel patch clamp experiments, mouse or human islets were dispersed with Accutase (Sigma) and cells were plated on sterilized glass shards. Recordings were performed according to (Krippeit-Drews et al., 2003). Gigaseals were obtained in extra cellular bath solution (in mM): 140 NaCl, 5 KCl, 1.2 MgCl_2_, 2.5 CaCl_2_, 0.5 Glucose, 10 HEPES, pH 7.4, adjusted with NaOH and clamped at −50mV before excision into inside-out configuration. Equilibrium solutions with K^+^ as the charge carrier were used for recording. The bath solution contained (in mM): 130 KCl, 2 CaCl_2_, 10 EGTA, 0.9 free Mg^2+^, 10 sucrose, 20 HEPES, pH 7.2 with KOH. The pipette solution contained (in mM): 10 sucrose, 130 KCl, 2 CaCl_2_, 10 EGTA, 20 HEPES, pH 7.2, adjusted with KOH. Intracellular recording electrodes made of borosilicate glass (Harvard Apparatus, Holliston, MA) with tip resistance of 3 MΩ (Sutter Instruments P-1000) were polished by microforge (Narishige MF-830) to the final tip resistance of 5-10 MΩ. After formation of gigaseal (>2.5 GΩ) and withdrawal of the pipette, excised inside-out configuration was established. A HEKA Instruments EPC10 patch-clamp amplifier was used for registration of current. Data was filtered online at 1kHz with Bessel filter and analyzed offline using Clampfit 10 software (Axon Instruments).

### Mouse islet GSIS

After incubation, equal numbers of islets were hand-picked and placed into a 12-channel BioRep device (75-100 islets/chamber) containing 2.7 mM glucose KRPH buffer (in mM: 140 NaCl, 4.7 KCl, 1.5 CaCl_2_, 1 NaH_2_PO4, 1 MgSO_4_, 2 NaHCO_3_, 5 HEPES, 1% fatty acid-free BSA, and 1 mM leucine as indicated; pH 7.4) with 100 µL Bio-Gel P-4 Media (Bio-Rad). Islets were equilibrated for 48 minutes, and then perifused in intervals based on the experimental conditions. Insulin and glucagon content and secretion was assessed by Insulin AlphaLISA (Perkin Elmer) with an alpha plate reader (TECAN Spark). After the perifusion, islets from each chamber were collected and incubated at room temperature for 15 minutes in 1 ml 0.1% Triton-X. Islets were then vortexed and frozen. DNA content was determined using the Quant-IT Pico Green dsDNA Assay Kit (Thermo Fisher Scientific).

### Respirometry

Islet oxygen consumption rates (OCR in pmoles/min) were measured with the Seahorse XF-24 Analyzer, and INS1 832/13 cells with the Seahorse XF-96 Analyzer (Agilent). Islets were recorded in DMEM (Sigma D5030, without glucose, glutamine, phenol red, FBS or sodium bicarbonate) with addition of 0.02 % bovine serum albumin, 2.5 mM glucose, and 5 μM PKa or vehicle (0.1% DMSO) (**Figure 5A**). 60-70 islets were deposited into each well of XFe24 plate, washed twice with the media, covered with an islet capture screen, and incubated for 1 hour before introduction to the Seahorse instrument. Baseline respiration was measured in 2.5 mM glucose. Islets were then sequentially exposed to 9 mM glucose, 1 μM oligomycin, 1 μM FCCP, and 5 μM rotenone as indicated. Oxygen consumption in INS1 832/13 cells was recorded in similar DMEM as the islets described above with additional 10 mM HEPES and pre-incubated with 2.5 mM glucose only (**Figures 5B, 6D**). Once in the instrument, baseline respiration was measured followed by stimulation with glucose as indicated. Cellular respiration was perturbed following the glucose addition by either 1 µM FCCP or 100 µM tolbutamide followed by 5 µM nimodipine. All cells were then finally exposed to a mixture of 5 µM rotenone and 10 µM antimycin A. For the studies with permeabilized INS1 832/13 cells in the presence of ADP and succinate, the cells were permeabilized using the XF Plasma Membrane Permeabilizer reagent based on the Agilent Technologies’ instructions (**Figures 6E, 6F**). The 1x MAS buffer (Agilent) with 10 mM succinate was used. The INS1 832/13 cells were treated with various ADP concentrations (62.5 µM, 125 µM, or 250 µM), 625 µM PEP, and 4 µM FCCP as indicated (**Figure 6E**). In the experiment for **Figure 6F** the cells were exposed to 125 µM ADP or 500 µM ADP (for the 0 µM PEP condition), then to various PEP concentrations (0 µM, 78 µM, 156 µM, 313 µM, or 625 µM), next to 1 mM ADP, and finally to antimycin A (10 µM) and rotenone (5 µM).

### PK Activity

The enzymatic assay of pyruvate kinase (EC 2.7.1.40) is from Sigma (Bergmeyer, H.U. et al.), with modifications for a 96-well plate format, and the use of a 1:2 PEP dilution curve starting at 2.5 mM or 5 mM PEP. Unless specified, 3 μM FBP was present.

### Human Islet Insulin Secretion Studies

Human islets from normal and Type 2 diabetic donors obtained from the University of Alberta Diabetes Institute, the University of Chicago Diabetes Research and Training Center, or the University of Minnesota Schulze Diabetes Institute were cultured in glutamine-free CMRL supplemented with 10 mM niacinamide and 16.7 μM zinc sulfate (Sigma), 1% ITS supplement (Corning), 5 mM sodium pyruvate, 1% Glutamax, 25 mM HEPES (American Bio), 10% HI FBS and antibiotics (10,000 units/ml penicillin and 10 mg/ml streptomycin). All media components were obtained from Invitrogen unless otherwise indicated. Human donor islets were cultured intact, then dispersed and re-aggregated as pseudo-islets for dynamic insulin secretion studies. Islets were dispersed with Accutase (Invitrogen) and the resulting cell suspension was seeded at 5000 cells per well of a 96-well V-bottom plate, lightly centrifuged (200*g*) and then incubated for 12-24 h at 37°C 5% CO_2_/95% air. GSIS perifusion studies were performed on an 8-channel BioRep device. The secretion buffer consisted of DMEM (Sigma D5030) supplemented with NaHCO_3_, 10 mM HEPES, 2 mM glutamine, 0.2% fatty acid-free BSA and 2.5 mM followed by 9 mM glucose in presence of 10 µM TEPP-46 or DMSO (0.1% final) adjusted to the same volume for the vehicle control. Additionally, dynamic GSIS studies were performed 24 h after the islets were plated into the 96 well plates following dispersion and re-aggregation. The human islet plates were washed and incubated at 37°C 5% CO_2_/95% air in standard KRB (in mM: 115 NaCl, 5 KCl, 24 NaHCO_3_, 2.2 CaCl_2_, 1 MgCl_2_) supplemented with 2.5 mM glucose, 2 mM glutamine, 24 mM HEPES and 0.25% BSA for 1.5 h. After the first incubation, human islet plates were then washed with glucose free KRB and incubated for 2 hours in KRB study media with 1, 2.5, 5, 7, 9, 11.2, 13 or 16.7 mM glucose in presence of 10 µM TEPP-46 or 0.1% DMSO. To induce glucolipotoxicity in normal donors, islet re-aggregates were incubated for 72 h under control or glucolipotoxic conditions (1% fatty acid free BSA alone or conjugated with 2:1 sodium oleate/sodium palmitate to a final concentration of 1 mM and 20 mM glucose), which were removed for the acute study. Supernatants were evaluated by insulin ELISA (ALPCO).

### ^13^C-Isotopic Labeling Studies

INS1 832/13 cells were cultured as monolayers in RPMI-1640 complete medium as previously described (Kibbey et al., 2007). ^13^C-Isotopic labeling studies were performed in DMEM medium (D5030, Sigma-Aldrich) supplemented with 9 mM glucose, 4 mM glutamine, 0.05 mM pyruvate, and 0.45 mM lactate. Cells were preincubated in this media for 2 h until a metabolic steady state was reached at which time unlabeled glucose was replaced with [U-^13^C_6_]glucose (Cambridge Isotope Laboratories). Following the addition of ^13^C-label, cells were quenched at t = 0, 5, 15, 30, 60, 120 and 240 min (*n* = 6 per time point). Cell quenching and sample preparation was performed as previously described (Alves et al., 2015; Patel et al., 2010).

### LC-MS/MS Analysis and modeling of ^13^C-labeled time courses

Metabolite concentrations and ^13^C-enrichments were determined by mass spectrometry using a SCIEX 5500 QTRAP equipped with a SelexION for differential mobility separation (DMS) as described previously (Alves et al., 2015). The integrated analysis of ^13^C labeling time courses was performed using a mathematical model of the TCA cycle similar to the one described previously (Alves et al., 2015). The model describes the transfer of label through the distal portion of glycolysis and the TCA cycle using [U-^13^C_3_]DHAP as a driving function. The label is distributed through all possible isotopomers for PEP, pyruvate, citrate, αKG, glutamate, succinate, malate and OAA and it is used to measure the flux through glycolysis, pyruvate kinase, pyruvate dehydrogenase, pyruvate carboxylase, citrate synthase, reversed isocitrate dehydrogenase and β-oxidation. Isotopomers were grouped in combination pools based on the number and/or position of labeled carbons and used to fit target data. Metabolic modeling and statistical analysis was performed using CWave software, version 4.0 (Mason et al., 2003) running in MATLAB. The flux values is shown as the least-square fit ± the standard deviation of the distribution of uncertainty from 100 Monte-Carlo simulations (Patel et al., 2010).

### Statistics

Data are expressed as means ± SE. Statistical significance was determined using one- or two-way ANOVA with Sidak multiple-comparisons test post hoc or Student’s t-test as appropriate. Differences were considered to be statistically significant at *P* < 0.05. Statistical calculations were performed with GraphPad Prism.

### Study approval

All procedures involving animals were approved by the Institutional Animal Care and Use Committees of the University of Wisconsin-Madison and the William S. Middleton Memorial Veterans Hospital, and followed the NIH Guide for the Care and Use of Laboratory Animals (8th ed. The National Academies Press. 2011.).

