## Supplemental Figures for "Pyruvate kinase controls signal strength in the insulin secretory pathway"

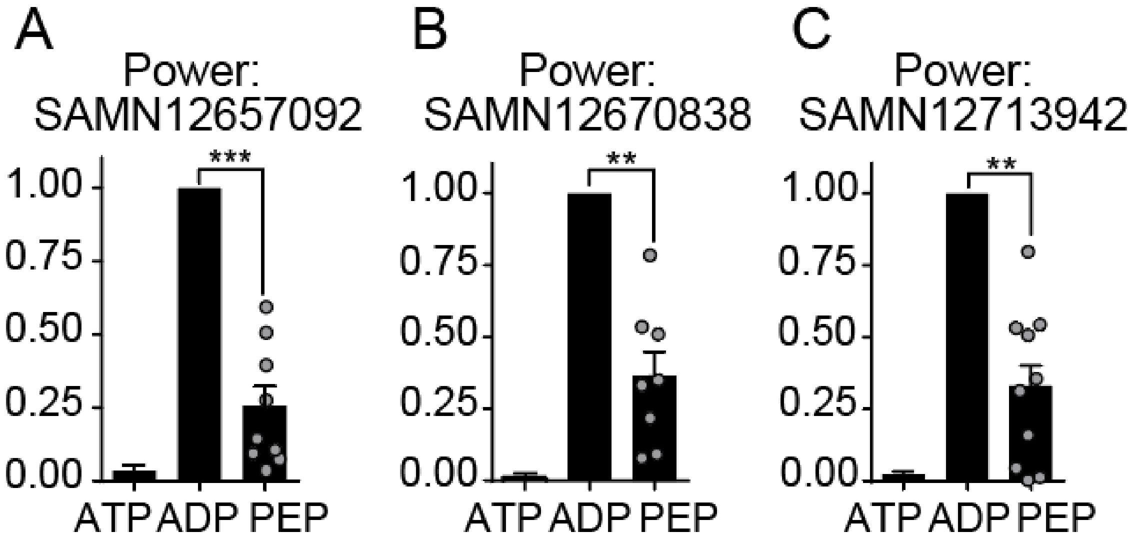

**Figure S1. Membrane-associated PK is sufficient to close $K_{ATP}$ channels in human islets, Related to Figure 1**

(A-C) Analysis of $K_{ATP}$ channel closure in terms of power for each human islet donor. Data are shown as mean ± SEM. $**p < 0.01$, $***p < 0.001$ by 1-way ANOVA.

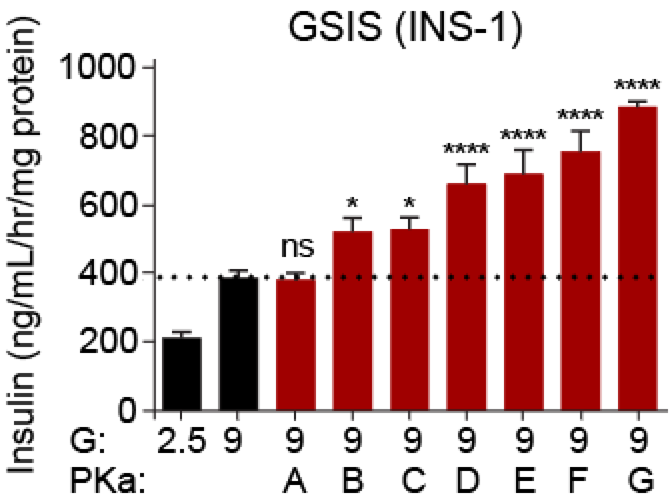

**Figure S2. PK activators stimulate insulin secretion in INS-1 cells, Related to Figure 2**
PK activators increased the average insulin secretion of INS-1 cells in a static incubation in 9 mM glucose (9G). A, 10 µM NIH NCGC185916-06 (Dasa-58); B, 10 µM NIH NCGC188799-02; C, 10 µM TEPP-46; D, 10 µM NIH NCGC186527-05; E, 10 µM NIH NCGC181801-02; F, 10 µM NIH NCGC188795-01; G, 10 µM NIH NCGC183333-05.
Data points represent the mean of 6 technical replicates for each experiment and are shown as mean ± SEM. *$p < 0.05$, ****$p < 0.0001$ by 1-way ANOVA.

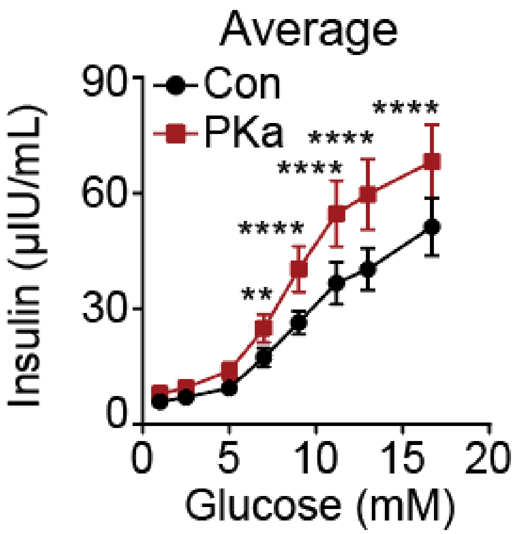

**Figure S3. PK activators enhance average insulin secretion from healthy human islets, Related to Figure 2**

PKa increased the average insulin secretion from all 10 donors from Figure 3 in static incubation assays.

Data points represent the mean of 4 technical replicates and are shown as mean ± SEM. **$p < 0.01$, ****$p < 0.0001$ by 1-way ANOVA.

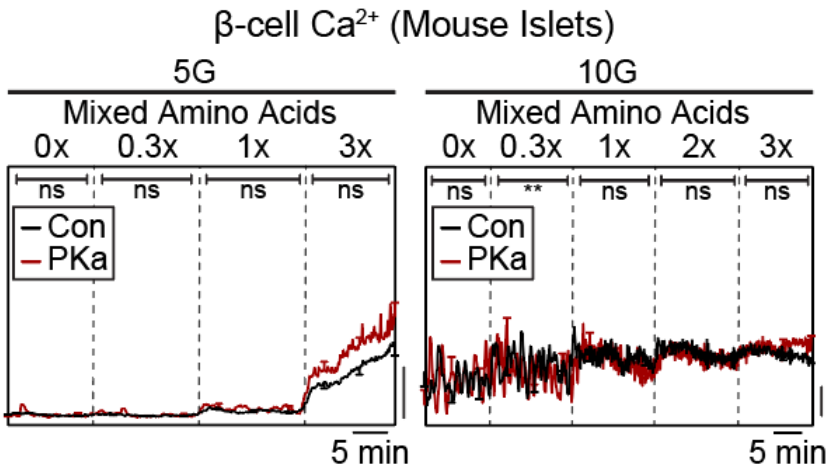

**Figure S4. The effect of PK activation on β-cell $Ca^{2+}$ is diminished at elevated glucose concentrations, related to Figure 4**

Representative average β-cell calcium in the absence or presence of PKa and in response to an amino acid ramp at 5 mM glucose (5G, left), and 10G (right) in mouse islets. (Left) PKa slightly increased β-cell calcium in 5G (Con, n = 20; PKa, n = 19). (Right) PKa did not alter the β-cell calcium response in 10G (Con, n = 14; PKa, n = 13).

Data are shown as mean ± SEM. **$p < 0.01$ by t-test.

| Donor | Age (years) | Sex | BMI | HbA1c % |
|---|---|---|---|---|
| SAMN12657092 | 42 | F | 32.7 | N/A |
| SAMN12670838 | 40 | M | 30.7 | N/A |
| SAMN12713942 | 41 | M | 23.5 | N/A |
| R073 | 74 | F | 28.3 | 5.4 |
| R034 | 76 | F | 23.7 | 5.9 |
| H1009 | 52 | M | 34.2 | N/A |
| R081 | 68 | M | 23.7 | 5.9 |
| R082 | 65 | F | 24.9 | 5.4 |
| R076 | 63 | F | 26.5 | N/A |
| H108 | 32 | F | 36.2 | 5.0 |
| CHI 7/29 | 30 | N/A | 30 | N/A |
| R075 | 27 | M | 26.2 | 5.4 |
| R066 | 44 | M | 32.2 | N/A |
| CHI R34 | 59 | M | 20.71 | N/A |
| ***AVG±SEM*** | 50.9±4.4 | | 28.11±1.25 | 5.60±0.17 |

**Table S1. Human Islet Donors, Related to Figures 1-4**
Summary characteristics of donors studied. BMI – Body Mass Index, HbA1c – glycated hemoglobin.

| Amino acid | Concentration at 1x (μM) |
|---|---|
| Alanine | 2100 |
| Glutamine | 600 |
| Glycine | 700 |
| Valine | 550 |
| Leucine | 500 |
| Serine | 350 |
| Arginine | 200 |
| Lysine | 218 |
| Threonine | 121 |

**Table S2. Physiological amino acid mixture concentrations, Related to Figure 4**
